# Carboxypeptidase activity drives L,D-transpeptidase essentiality during vegetative growth and sporulation in *Clostridioides difficile*

**DOI:** 10.64898/2026.06.30.735746

**Authors:** Kevin W. Bollinger, Ute Müh, Paige B. Brannen, David L. Popham, David S. Weiss, Craig D. Ellermeier

## Abstract

In most bacteria, peptidoglycan contains mainly 4-3 crosslinks formed by penicillin-binding proteins (PBPs). But in the opportunistic pathogen *Clostridioides difficile*, 70% of the crosslinks are 3-3 crosslinks formed by L,D-transpeptidases (LDTs), and LDTs are essential for viability. PBPs and LDTs use different acyl donors for crosslinking; PBPs require a pentapeptide, while LDTs require a tetrapeptide. Here, we determined the source of the tetrapeptides in *C. difficile* and investigated the consequences of reengineering PG crosslinking from predominantly 3-3 to exclusively 4-3. We found that two D-alanyl-D-alanine carboxypeptidases (DD-CPase), DacA and DacC, supply LDTs with tetrapeptides during vegetative growth. Deleting these enzymes was sufficient to bypass the normal requirement for LDTs. The resulting mutant (Δ*dacAC* Δ*ldt*) was remarkably healthy despite the absence of 3-3 crosslinks. Its only major phenotypic defect was a 3- to 4-log decrease in sporulation, which could, however, be overcome by deleting a third DD-CPase, *dacB*. These findings fill gaps in our understanding of the pathway for LD-transpeptidation in *C. difficile* and imply that LDTs are not essential components of the elongasome or divisome, both of which function well in the complete absence of LDTs, provided there is sufficient pentapeptide to sustain crosslinking by PBPs. Thus, LDTs are essential for viability because *C. difficile* has intrinsically high levels of DD-CPase activity. Finally, we propose a model for how PBPs and LDTs work together during PG synthesis. In this model, PBPs construct a sparsely crosslinked PG sacculus that is subsequently strengthened with crosslinks introduced by LDTs.

**Importance:** Synthesis of peptidoglycan (PG) in the opportunistic gut pathogen *Clostridioides difficile* depends mostly on 3-3 crosslinks created by L,D-transpeptidases (LDTs). Here we converted *C. difficile* from an organism that relies primarily on 3-3 crosslinks made by LDTs to one that relies exclusively on 4-3 crosslinks made by an alternative family of crosslinking enzymes, the penicillin-binding proteins (PBPs). Our findings explain why 3-3 crosslinking predominates over 4-3 crosslinking, why *C. difficile* does not compensate for loss of LDTs by increasing PBP activity, and suggest a model for how PBPs and LDTs work together in this problematic pathogen.

## Introduction

Peptidoglycan (PG) assembly is the target of many highly successful antibiotics because it is unique to bacteria and essential for protecting them from lysis due to turgor pressure [1]. PG encases the cytoplasmic membrane in a net-like sacculus composed of glycan strands held together by peptide crosslinks [2, 3]. In model organisms such as *Escherichia coli*, over 90% of the crosslinks are formed by DD-transpeptidases commonly referred to as penicillin-binding proteins (PBPs) [4]. PBPs are the lethal target of penicillin and other β-lactam antibiotics, which irreversibly acylate a serine in the catalytic site [5]. The remaining ∼10% of the crosslinks are formed by LD-transpeptidases (LDTs), which are not typically essential for viability and are only inactivated by one class of β-lactams, the penems [4, 6, 7].

An exception to this paradigm is the opportunistic gut pathogen *Clostridioides difficile*. In *C. difficile*, about 70% of the crosslinks are formed by LDTs and about 30% by PBPs, and both types of crosslinking enzymes are essential for viability [8–13]. The only other bacterium in which LDTs are known to be essential for viability is the plant pathogen *Agrobacterium tumefaciens* [6]. The nearly unique essentiality of LDTs in *C. difficile* suggests they might be exploited as targets for a “silver bullet” antibiotic that does not disrupt the normal gut microbiota, which is important for keeping *C. difficile* in check [12].

PBPs and LDTs have different substrate requirements and form chemically distinct crosslinks. PBPs utilize a pentapeptide acyl donor substrate to create a 4-3 crosslink between the 4^th^ position D-alanine and 3^rd^ position meso-diaminopimelic acid (mDAP) from peptide sidechains on adjacent glycan strands [5]. In contrast, LDTs require a tetrapeptide acyl donor and create a 3-3 crosslink between the third position mDAP residues of two PG peptide sidechains [7, 14]. In a previous report, we observed that depletion of LDTs in *C. difficile* results in loss of 3-3 crosslinks with no compensating increase in 4-3 crosslinks, leading to loss of PG integrity, bulging and cell lysis [12].

PG is synthesized from a lipid-linked disaccharide-pentapeptide precursor known as lipid II. Glycosyltransferases polymerize the disaccharides to create nascent glycan chains that are then crosslinked by PBP transpeptidases [3]. The tetrapeptide substrate required by LDTs as an acyl donor can be generated in two ways. The first and most common is the removal of the last D-Ala residue from D-Ala^4^-D-Ala^5^ termini of pentapeptide precursors by enzymes known as D-alanyl-D-alanine carboxypeptidases (DD-CPases) encoded by so-called *dac* genes [5, 14, 15]. The second is cleavage of existing 4-3 crosslinks by PG endopeptidases [16]. A recent report implicated one or more DD-CPases as the source of tetrapeptides crosslinked by LDTs in *C. difficile*, but the relevant enzymes have not been identified [17].

Here we address a series of interrelated questions concerning the relationship between 3-3 and 4-3 crosslinking in *C. difficile*. What is the source of the tetrapeptide substrate required by LDTs? Why doesn’t *C. difficile* respond to loss of 3-3 crosslinks by increasing 4-3 crosslinks? Can PG synthesis be engineered in *C. difficile* to create an exclusively 4-3 crosslinked PG sacculus? If so, what are the phenotypic consequences?

## Results

### Loss of *dacA* and *dacC* circumvents LDT essentiality

To identify the relevant DD-CPase(s), we took advantage of an LDT-depletion strain described previously [12]. *C. difficile* has a total of five LDTs, three of which are canonical YkuD-family enzymes [8, 18, 19], while the other two are of a newly discovered family that has a VanW catalytic domain [12]. In the depletion strain, the genes for four LDTs have been deleted and expression of the fifth is controlled by the P*_tet_* promoter (Δ*ldt1-4 P_tet_::ldt5*). Growth of the depletion strain in the absence of the inducer anhydrotetracycline (aTet) results in loss of 3-3 crosslinks, bulging and cell lysis [12]. We hypothesized that PBP-mediated 4-3 crosslinking cannot rescue viability in the absence of LDTs because DD-CPases convert pentapeptides to tetrapeptides, leaving the PBPs without suitable crosslinking substrates. This hypothesis predicts that deletion of DD-CPase genes will improve 4-3 crosslinking and might even bypass the requirement for LDTs altogether.

There are seven predicted DD-CPases in the R20291 genome, of which four rely on a serine nucleophile for catalysis and three rely on a metal such as Zn^++^ (Table 1). The serine-type DD-CPases are also called low molecular weight PBPs. In contrast to high molecular weight PBPs, these enzymes hydrolyze rather than crosslink the acyl donor peptide substrate. Nevertheless, low molecular weight PBPs are susceptible to inactivation by β-lactams. Based on RNA-sequencing, only three of the DD-CPases are expressed to any appreciable extent during vegetative growth [12]. These are the serine-type enzymes encoded by *cdr20291_0441* (*dacA*) and *cdr20291_2390* (*dacC*), and the metallo-type enzyme encoded by *cdr20291_2396* (*dacD*). Each of these has a predicted N-terminal Sec system signal peptide ending in a cleavage site for Type I signal peptidase, implying they are secreted across the cytoplasmic membrane.

**Table 1:** List of putative D-alanyl-D-alanine carboxypeptidases in *C. difficile*.

| Gene Locus | Gene Name | RPKM* | Carboxypeptidase-type |
| --- | --- | --- | --- |
| CDR20291_0441 | *dacA* | 2023 | Serine-type |
| CDR20291_2048 | *dacB* | 16 | Serine-type |
| CDR20291_2390 | *dacC* | 4575 | Serine-type |
| CDR20291_1131 | *dacF* | 68 | Serine-type |
| CDR20291_2396 | *dacD* | 357 | Metallo-type |
| CDR20291_3439 | *dacJ* | 91 | Metallo-type |
| CDR20291_1525 | *vanY* | 64 | Metallo-type |

\*Reads per kilobase of transcript per million mapped reads

To determine if eliminating these DD-CPases can bypass the essentiality of LDTs, we used CRISPR mutagenesis to delete the three *dac* genes alone and in combination in the LDT-depletion background. Viability was assessed by spot titer assays on TY plates with or without aTet.

Deletion of *dacA* or *dacC* slightly increased viability in the absence of aTet, while a Δ*dacAC* double deletion resulted in wild type (WT) levels of viability (Fig. 1). In contrast, deletion of *dacD* alone or in combination with *dacA* and/or *dacC* had no effect (Fig. 1). Expression of *dacA* or *dacC* from a plasmid in the Δ*dacAC*Δ*ldt1-4 P_tet_::ldt5* background caused a severe reduction in viability, confirming that loss of *dacA* and *dacC* was responsible for suppressing the essentiality of LDTs (Fig. S1). Thus, the two serine-type DD-CPases that are expressed during vegetative growth are responsible for making LDTs essential in *C. difficile*, likely because they convert PG pentapeptides to tetrapeptides that LDTs can utilize but PBPs cannot. Moreover, because both DD-CPases are predicted to be exported, conversion of pentapeptides to tetrapeptides presumably happens outside of the cell, which is consistent with the finding that low doses of β-lactams skew crosslinking from 3-3 to 4-3 [17].

**FIG 1:**
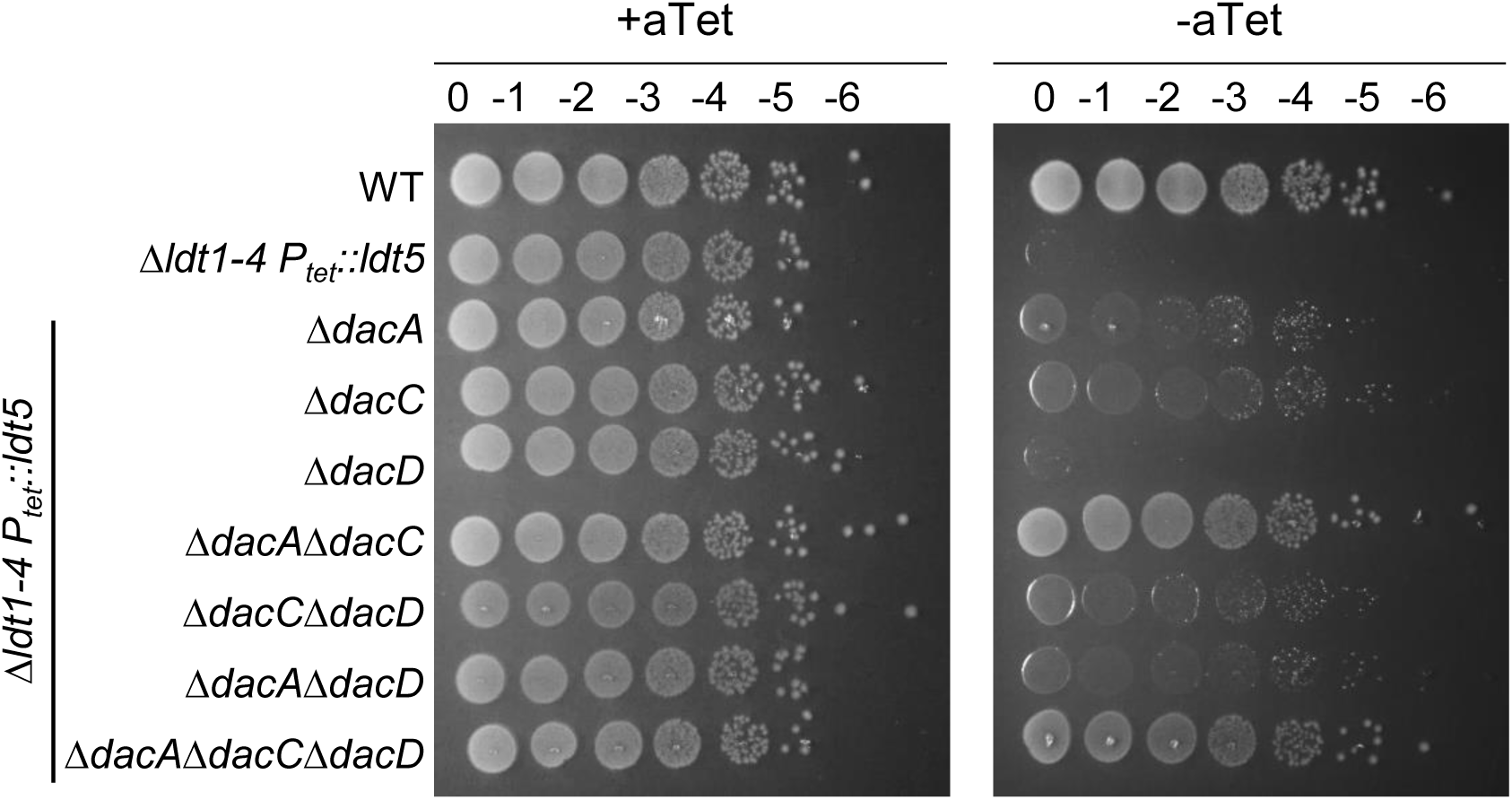
Loss of DD-carboxypeptidase activity increases viability of the LDT depletion strain. Spot titer assays of LDT depletion strain with deletions of various carboxypeptidase genes. Tenfold serial dilutions were spotted on TY plates with or without 25 ng/mL anhydrous tetracycline (aTet). Plates were photographed after incubation for 20 h. Images are a representative of three experiments.

### Growth rate, morphology and toxin secretion are largely normal in the absence of DD-CPases and LDTs

To rule out the possibility that viability of the Δ*dacAC*Δ*ldt1-4 P_tet_::ldt5* strain in the absence of aTet depends on leaky expression of *P_tet_*::*ldt5*, we used CRISPR to delete *ldt5*. The resulting Δ*dacAC*Δ*ldt1-5* mutant was confirmed by whole genome sequencing and will be referred to henceforth as Δ*dacAC*Δ*ldt* for brevity. There was no significant difference in the growth rate between WT, Δ*dacAC*, and Δ*dacAC*Δ*ldt* in TY broth (Fig. 2A). Microscopy of cells harvested in exponential growth revealed only minor changes in morphology in the Δ*dacAC*Δ*ldt* mutant. Although cell length was the same as WT, the Δ*dacAC*Δ*ldt* mutant was ∼10% thinner (0.81 versus 0.75 μm) and ∼17% of the cells exhibited a more curved or irregular contour than WT, which grew as straight rods (Fig. 2B-E). These changes in shape were due to the lack of LDTs rather than lack of the two DD-CPases because the morphology of the Δ*dacAC* mutant was similar to WT in all respects (Fig. 2B-E).

**FIG 2:**
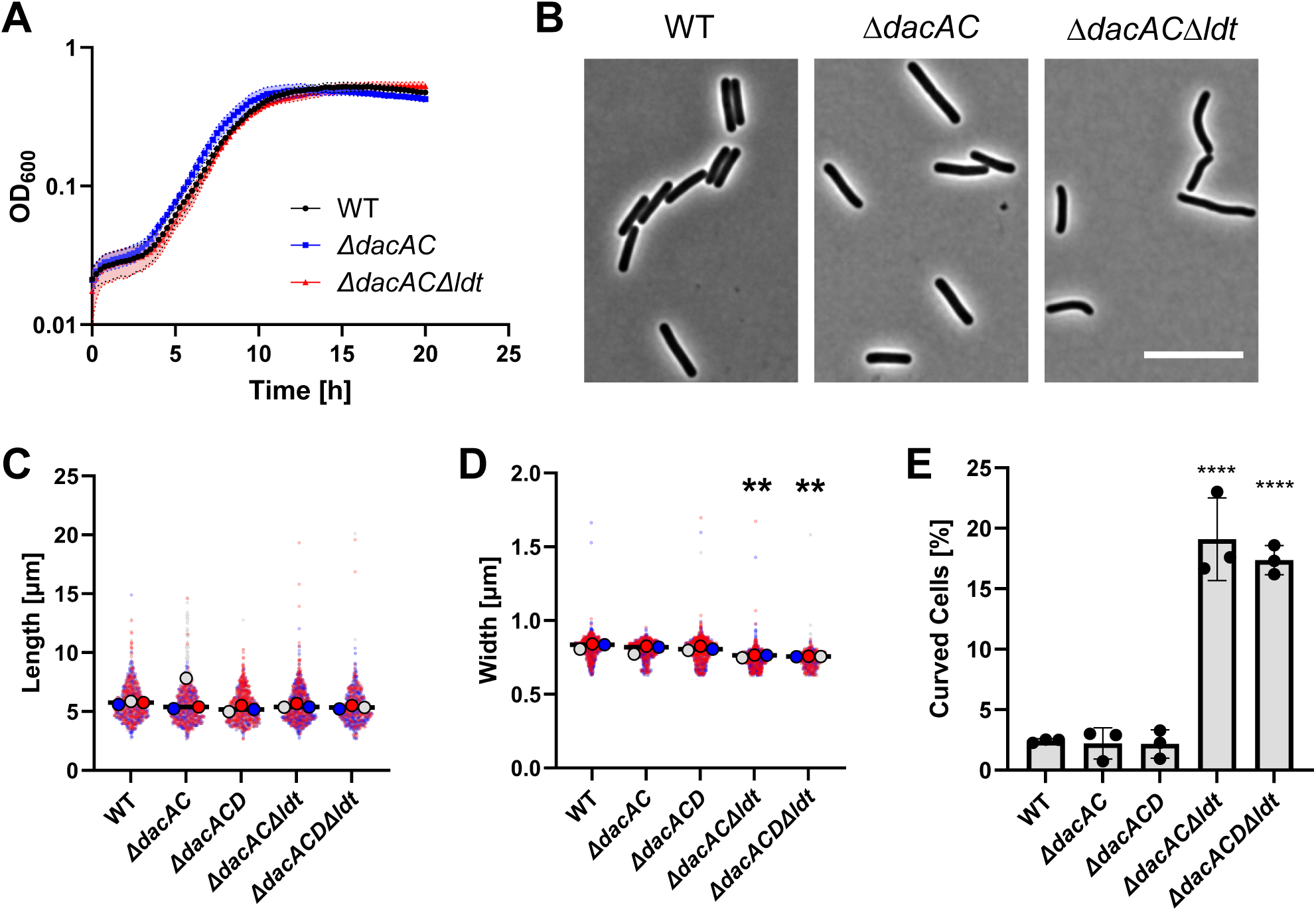
Growth and morphology of an LDT null mutant. (A) Growth curves of WT, Δ*dacAC*, and Δ*dacAC*Δ*ldt* in TY broth. Data are graphed as mean +/- SD of four biological replicates on different days. (B) Phase contrast microscopy of cells sampled during mid-log growth, about 5.5 hours after subculture. Images are representative of three biological replicates. Size bar = 10 µm. (C-E) Quantification of cell length, width, and sinuosity. Cell length and width are graphed as superplots pooling data from three biological replicates distinguished by different colors. Small dots depict measurements of individual cells. Large dots are the means of each replicate. Sinuosity is a measure of curvature and is graphed as the percentage of cells with a sinuosity of >1.03 plotted as the mean and SD of three biological replicates. For C-E ≥115 cells were measured per experiment. Statistical significance was calculated for D and E using a one-way ANOVA followed by a Tukey’s test comparing WT to the other strains based on the averages for each biological replicate. ** = p<0.01; **** = p<0.0001.

To ask whether DacD makes subtle contributions to pentapeptide processing that were not apparent from the viability assays in the LDT depletion background, we constructed a Δ*dacACD*Δ*ldt* deletion mutant. Its viability and morphology were indistinguishable from the Δ*dacAC*Δ*ldt* parent (Fig. 1; Fig. 2C-E). Thus, we have no evidence that DacD contributes to the production of tetrapeptide substrates for the LDTs, although it might do so under conditions we have not tested.

The severe intestinal inflammation associated with *C. difficile* infections is caused by two toxins, TcdA and TcdB, that glucosylate Rho family GTPases [20]. Both toxins are secreted by an atypical mechanism involving the holin-like protein TcdE [20, 21]. In *Salmonella* Typhi, secretion of the typhoid toxin is facilitated by an LDT [22, 23], so we asked whether the same might be true of TcdA and TcdB in *C. difficile*. However, similar amounts of both toxins were detected by immunoblot in cell-free spent media from cultures of WT, Δ*dacAC*, and Δ*dacAC*Δ*ldt* (Fig. S2). We conclude that DD-CPases and LDTs are not required for the secretion of TcdA or TcdB.

The experiments described above used cells harvested at a single timepoint, 5.5 hours after sub-culture (OD_600_ ∼0.6). To monitor morphology during active growth, we turned to time-lapse microscopy of cells immobilized on an agarose pad in a Gene Frame device to maintain anaerobiosis as described by Shen and coworkers [24]. Images were captured every 5 min for up to 6 hours (Fig. 3, videos S1-S4). WT grew actively and without signs of lysis, but the LDT depletion strain (Δ*ldt1-4 P_tet_::ldt5*) lost rod shape, developed bulges and eventually lysed. These defects are consistent with loss of PG integrity due to the loss of 3-3 crosslinks as we reported previously [12]. The Δ*dacAC* strain grew as normal rod-shaped cells. Finally, the Δ*dacAC*Δ*ldt* strain proliferated well and without any signs of lysis, although the cells were somewhat curved compared to WT. Overall, time-lapse microscopy confirmed that (i) the Δ*dacAC*Δ*ldt* strain grows well albeit with modest morphological defects and (ii) the defects are due to loss of LDTs rather than loss of *dacAC*.

**FIG 3:**
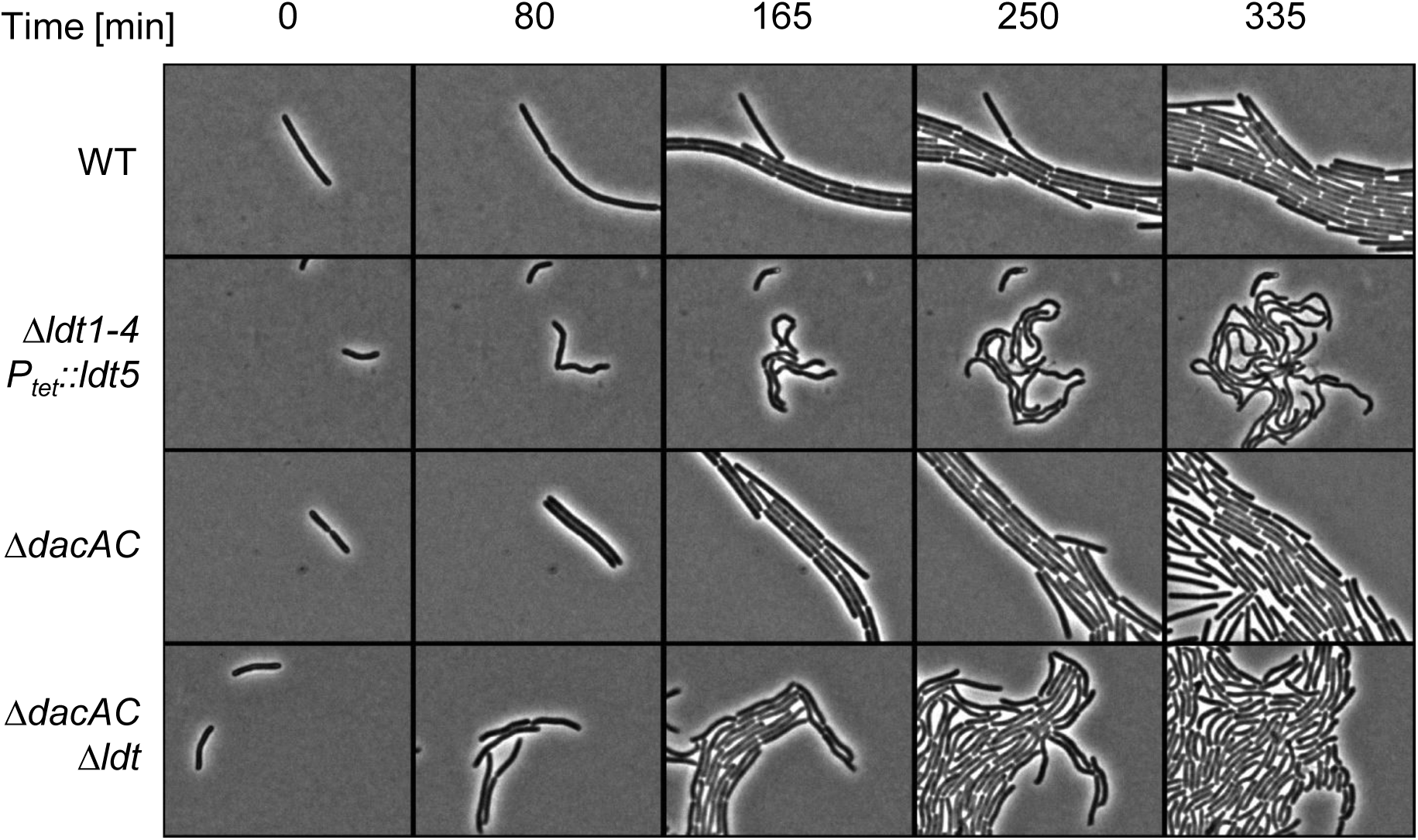
Time-lapse microscopy reveals deletion of LDTs in Δ*dacAC* background causes slight morphological defects. Overnight cultures were subcultured 1:100 into TY. After one hour, 1 µL samples were spotted onto an anaerobic agarose pad, sealed, and transferred to a microscope with a heated stage. Phase contrast images were captured every five minutes for about six hours. Images are representative of at least three experimental replicates done on different days.

### Loss of DD-CPases and LDTs shifts crosslinking from 3-3 to 4-3

To determine the impact of deleting genes for DD-CPases and LDTs on the nature and abundance of PG crosslinks, we analyzed muropeptides from WT, the Δ*dacAC* double mutant, and the Δ*dacAC*Δ*ldt* septuple mutant. Muropeptides were separated by reverse-phase high-pressure liquid chromatography (RP-HPLC) and detected by A_206_ as they eluted from the column. Peaks were collected from the Δ*dacAC* double mutant to determine muropeptide structures by mass spectrometry. Representative RP-HPLC elution profiles are shown in Fig. 4. The deduced structures of 16 muropeptides are presented in Table 2, along with their abundance as a percentage of total analyzed muropeptides.

**FIG 4:**
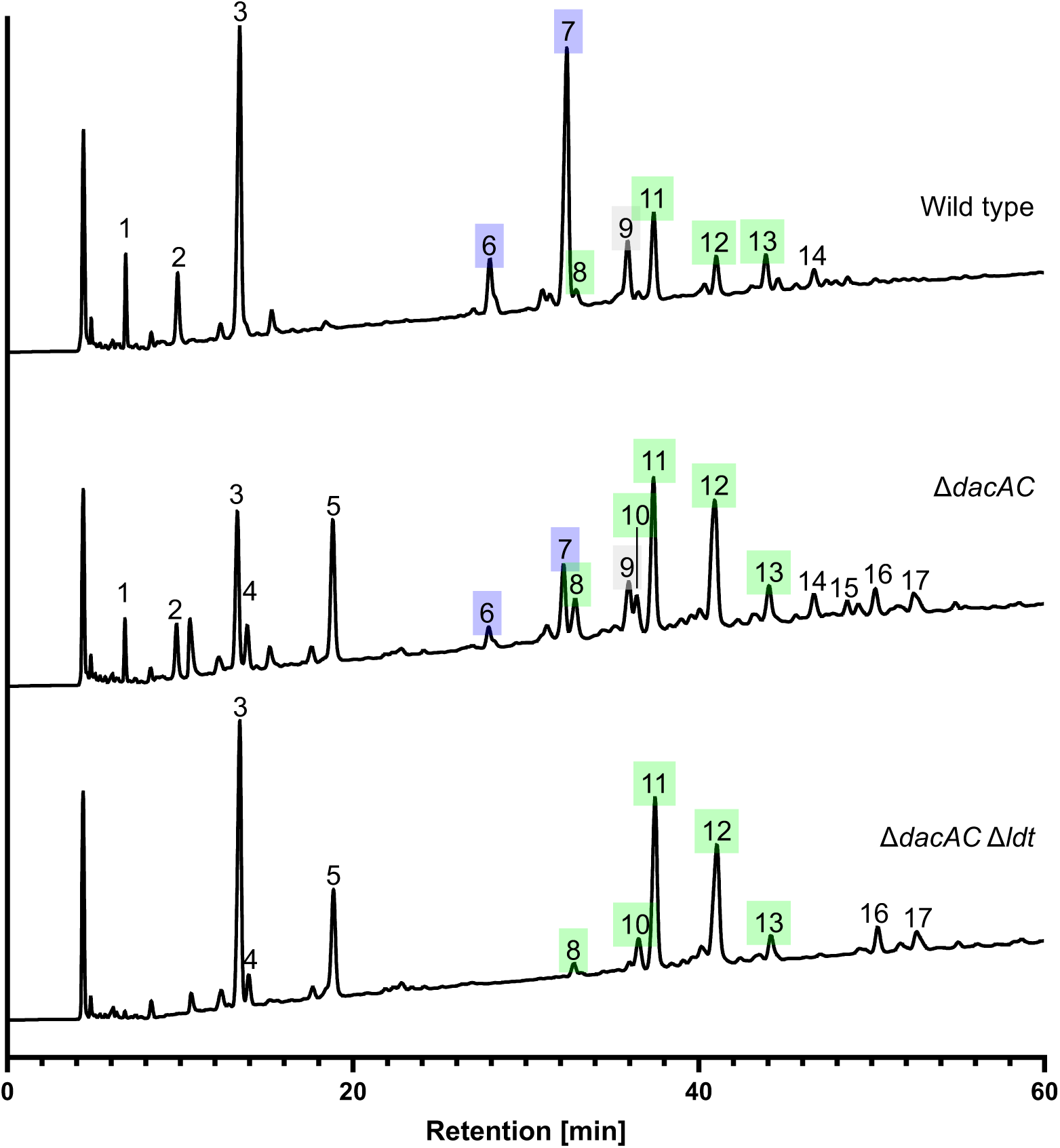
The absence of LDTs in the Δ*dacAC* background prevents the formation of 3-3 crosslinks in vegetative peptidoglycan. Vegetative PG was isolated from the indicated strains, digested with mutanolysin and reduced. The resulting muropeptides were separated by reverse-phase HPLC, detected as they eluted from the column by A_206_ and identified by mass spectrometry. Peak identification and quantitation are summarized in Table 2. Peak 1 is not a muropeptide. Peak numbers of dimeric species are highlighted based on crosslinking (blue: 3-3 and green: 4-3). Note that peak 9 (grey) is a mixture of 3-3 and 4-3 crosslinked species. Chromatograms are representative of three biological replicates.

**Table 2.**
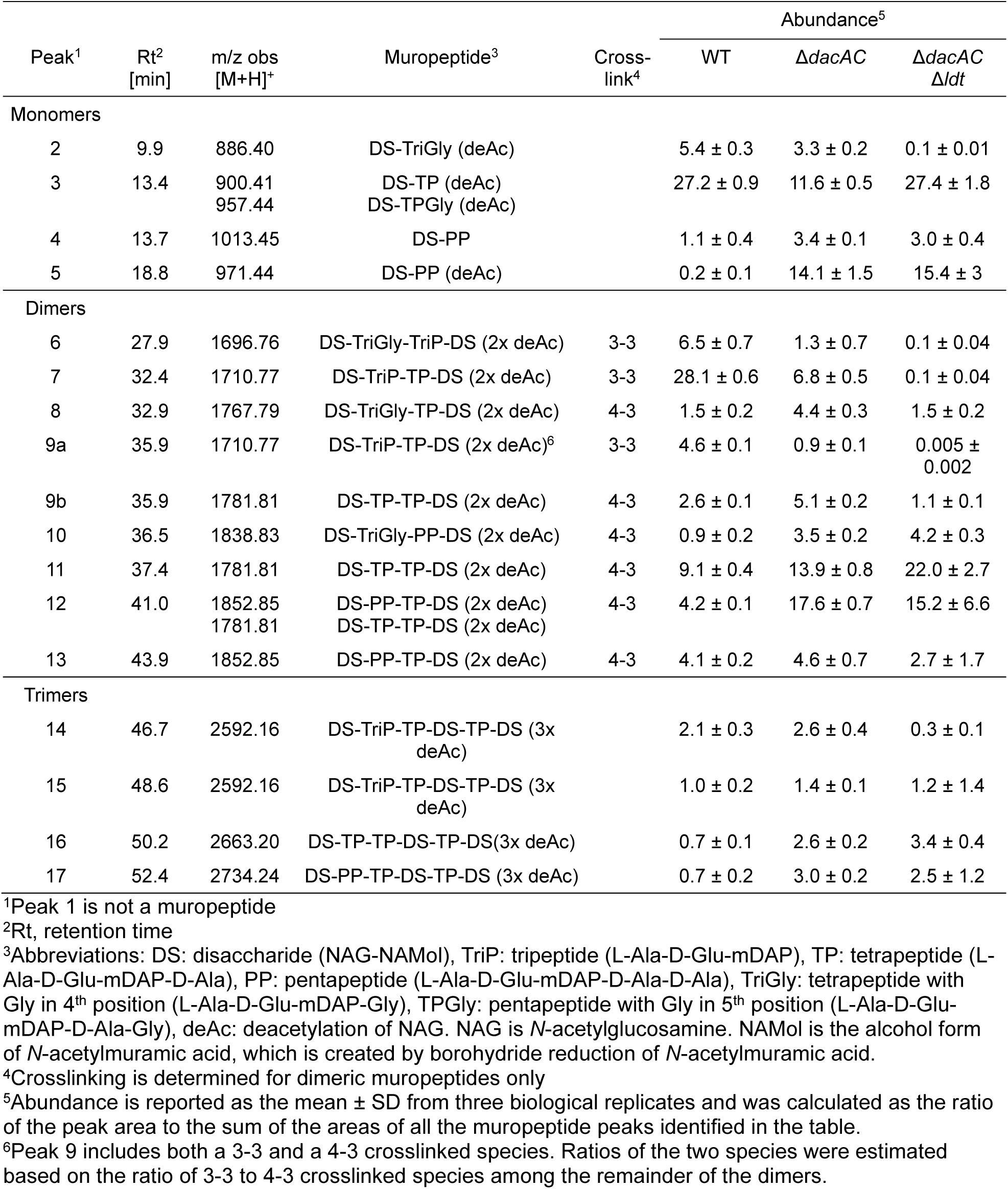
Muropeptide quantitation of vegetative peptidoglycan.

The muropeptide profile from WT revealed the presence of PG monomers, dimers and trimers. The abundance of these species as a percentage of total muropeptides was 34% monomers, 62% dimers, and 4% trimers, similar to previous reports [8, 12, 19]. Of the dimers, 64% were 3-3 crosslinked (peaks 6, 7, and 9a) and 36% were 4-3 crosslinked (peaks 8, 9b, 10-13). In the Δ*dacAC* double mutant, the relative abundance of monomers, dimers and trimers was essentially unchanged but there was a drastic increase in monomers with pentapeptide sidechains (peaks 4 and 5) at the expense of tetrapeptides (peaks 2 and 3). Moreover, the relative abundance of 3-3 to 4-3 crosslinked dimers flipped, with 4-3 now comprising 84% of the dimers. This shift indicates PBP activity is substrate-limited in WT and the enzymes primarily responsible for the conversion of pentapeptides to tetrapeptides are DacA and DacC. The remaining 3-3 crosslinks present in the Δ*dacAC* mutant could be due to LDT-mediated crosslinking of tetrapeptides generated when endopeptidases cleave 4-3 crosslinks during PG remodeling and maturation. Finally, in the Δ*dacAC*Δ*ldt* strain, only 4-3 crosslinked dimers were unambiguously detected. Although Table 2 reports 3-3 crosslinked dimers at a relative abundance of 0.2%, this is below our detection limit and reflects assignment of subtle irregularities in the baseline A_206_ elution profile to 3-3 dimers based on retention time. We conclude that, within the detection limits of our methods, the Δ*dacAC*Δ*ldt* strain has an exclusively 4-3 crosslinked PG sacculus.

### LDTs confer modest resistance to glycopeptide antibiotics but not to β-lactams

It is unclear whether heavy reliance on 3-3 crosslinking by LDTs provides *C. difficile* with any protection against β-lactams or vancomycin, which is the recommended treatment for most *C. difficile* infections [25, 26]. On one hand, there is precedent for such resistance in other organisms [15, 16, 27–29]. Resistance also makes sense biochemically because most β-lactams do not bind to LDTs and vancomycin does not bind to the tetrapeptides that LDTs use as acyl donors for crosslink formation. On the other hand, β-lactams and vancomycin prevent crosslinking by PBPs, two of which are essential for viability in *C. difficile* despite the high abundance of 3-3 crosslinks [9–11]. Moreover, β-lactams and/or vancomycin might inhibit LD-transpeptidation indirectly by preventing DD-CPases from generating tetrapeptides. Indeed, this was recently proposed to be the case for two classes of β-lactams, the cephalosporins and carbapenems, which skewed crosslinking ratios in a way that implies they inactivate the DD-CPases at a lower concentration than required to inactivate crosslinking by PBP1 and PBP2 [17].

Based on these considerations, we asked what role, if any, the LDTs play in the susceptibility of *C. difficile* to several cell wall-targeting antibiotics: Three β-lactams from different classes (ampicillin, cefoxitin, and meropenem), bacitracin, daptomycin, and four glycopeptides (vancomycin, teicoplanin, dalbavancin, and oritavancin). The DNA gyrase inhibitor novobiocin served as a control. A shift in MIC was only observed for the glycopeptides (Fig. 5A). The Δ*dacAC*Δ*ldt* mutant was 2-fold more sensitive to vancomycin and 4- to 8-fold more sensitive to the other three glycopeptides, all of which have a lipophilic sidechain that anchors them in the membrane. Follow-up experiments using plasmid-based expression of individual LDTs revealed that Ldt1, Ldt4, or Ldt5 are sufficient to restore oritavancin resistance back to WT levels (Fig. 5B). These are the same three LDTs we previously demonstrated are sufficient for viability of *C. difficile* [12]. Expression of Ldt2 also somewhat improved resistance, but this change was not statistically significant (Fig. 5B). In summary, LDTs modestly improve resistance to glycopeptides, especially the lipoglycopeptides, but not to β-lactams.

**FIG 5:**
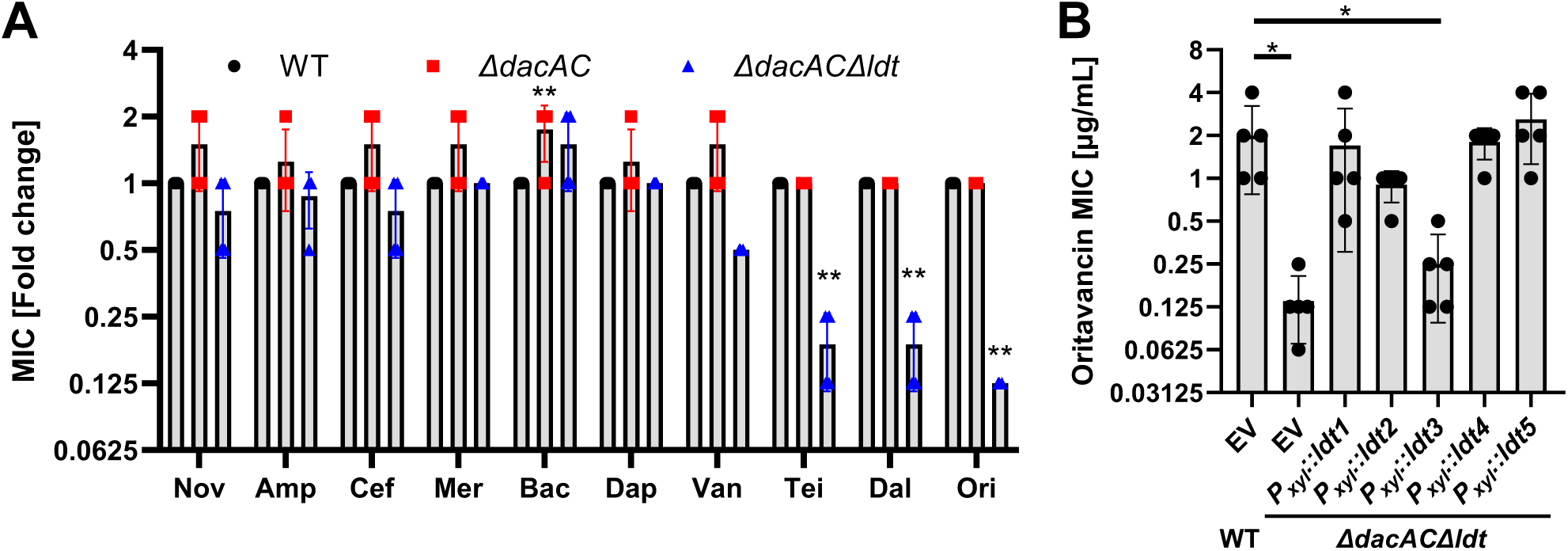
Δ*dacAC*Δ*ldt* is more sensitive to certain glycopeptides. (A) Minimum inhibitory concentration (MIC) assays were performed on the DNA gyrase inhibitor Novobiocin (Nov) and the following cell wall-targeting antibiotics: Ampicillin (Amp), Cefoxitin (Cef), Meropenem (Mer), Bacitracin (Bac), Daptomycin (Dap), Vancomycin (Van), Teicoplanin (Tei), Dalbavancin (Dal), and Oritavancin (Ori). MICs are plotted as the fold-change compared to wild-type based on four biological replicates. Bars, lines and colored symbols depict the mean, SD and individual measurements, respectively. The MICs determined for WT were: Nov: 18.75 µg/mL; Amp: 2.73 µg/mL; Cef: 187.5 µg/mL; Mer: 3.125 µg/mL; Bac: 312.5 µg/mL; Dap: 1.56 µg/mL; Van: 0.78 µg/mL; Tei: 0.3125 µg/mL; Dal: 0.156 µg/mL; Ori: 1 µg/mL. Statistical significance was calculated using a one-way ANOVA followed by a Tukey’s test. **=p<0.01. (B) MIC assays to test the effect of producing single LDTs from a P*_xyl_* expression vector on oritavancin resistance. EV = empty vector. Statistical significance was calculated using one-way ANOVA followed by a Dunnett’s test comparing WT EV to each of the other strains. * = p<0.05.

### LDTs are required for sporulation even in the absence of DacA and DacC

*C. difficile* pathogenesis relies on the formation of metabolically inert endospores that mediate transmission between hosts and facilitate recolonization of the colon after a course of antibiotic treatment is complete [30, 31]. Sporulation requires synthesis of two chemically distinct layers of PG [32–34]. One of these is the germ cell wall, which resembles vegetative PG and sits immediately outside of the inner spore membrane. The other is the spore cortex, a thick layer of highly modified and sparsely crosslinked PG located outside of the germ PG. Crosslinking of cortex PG is not well characterized in *C. difficile*, but the available evidence indicates the presence of both 3-3 and 4-3 crosslinks [17, 35, 36]. Cortex synthesis requires a dedicated PG synthase composed of the glycosyltransferase SpoVE and the monofunctional transpeptidase SpoVD, a PBP that makes 4-3 crosslinks [11, 37, 38]. No individual LDT is required for sporulation [9], but functional redundancy might mask their contributions. The requirement of at least one LDT for vegetative growth has made it impossible until now to ask whether LDTs are also essential for sporulation.

We plated WT, Δ*dacAC*, and Δ*dacAC*Δ*ldt* on sporulation media and quantified the number of ethanol resistant spores after 24 hours. High numbers of spores were recovered for both WT and Δ*dacAC*, but sporulation was down about 6000-fold for Δ*dacAC*Δ*ldt* (Fig. 6A). Expression of Ldt1, Ldt4, or Ldt5, but not Ldt2 or Ldt3, was sufficient to rescue sporulation of the Δ*dacAC*Δ*ldt* mutant to WT levels (Fig. 6B). The LDT requirement was confirmed by phase-contrast microscopy, which revealed numerous phase-bright spores in samples of WT and Δ*dacAC*, but not in Δ*dacAC*Δ*ldt*, where instead we observed mostly phase-dark bodies that could be cell debris or defective spores that cannot survive the ethanol treatment (Fig. 6C). To distinguish between these possibilities, we used a fusion of a codon optimized mCherry red fluorescent protein (RFP) to the promoter for *spoIIR*, a gene expressed in the forespore under SigF control (*P_spoIIR-_rfp*) [39–41]. As expected, microscopy of the WT *P_spoIIR-_rfp* reporter strain revealed numerous red fluorescent forespores (Fig. 7). In the Δ*dacAC*Δ*ldt P_spoIIR-_rfp* reporter strain, we observed red fluorescence in forespores and in some of the phase-dark bodies, indicating that SigF was activated despite the absence of mature spores (Fig. 7). Collectively, these findings demonstrate that LDTs are required for sporulation, deletion of *dacA* and *dacC* is not sufficient to bypass this requirement, and sporulation in the Δ*dacAC*Δ*ldt* mutant goes awry after asymmetric division.

**FIG 6:**
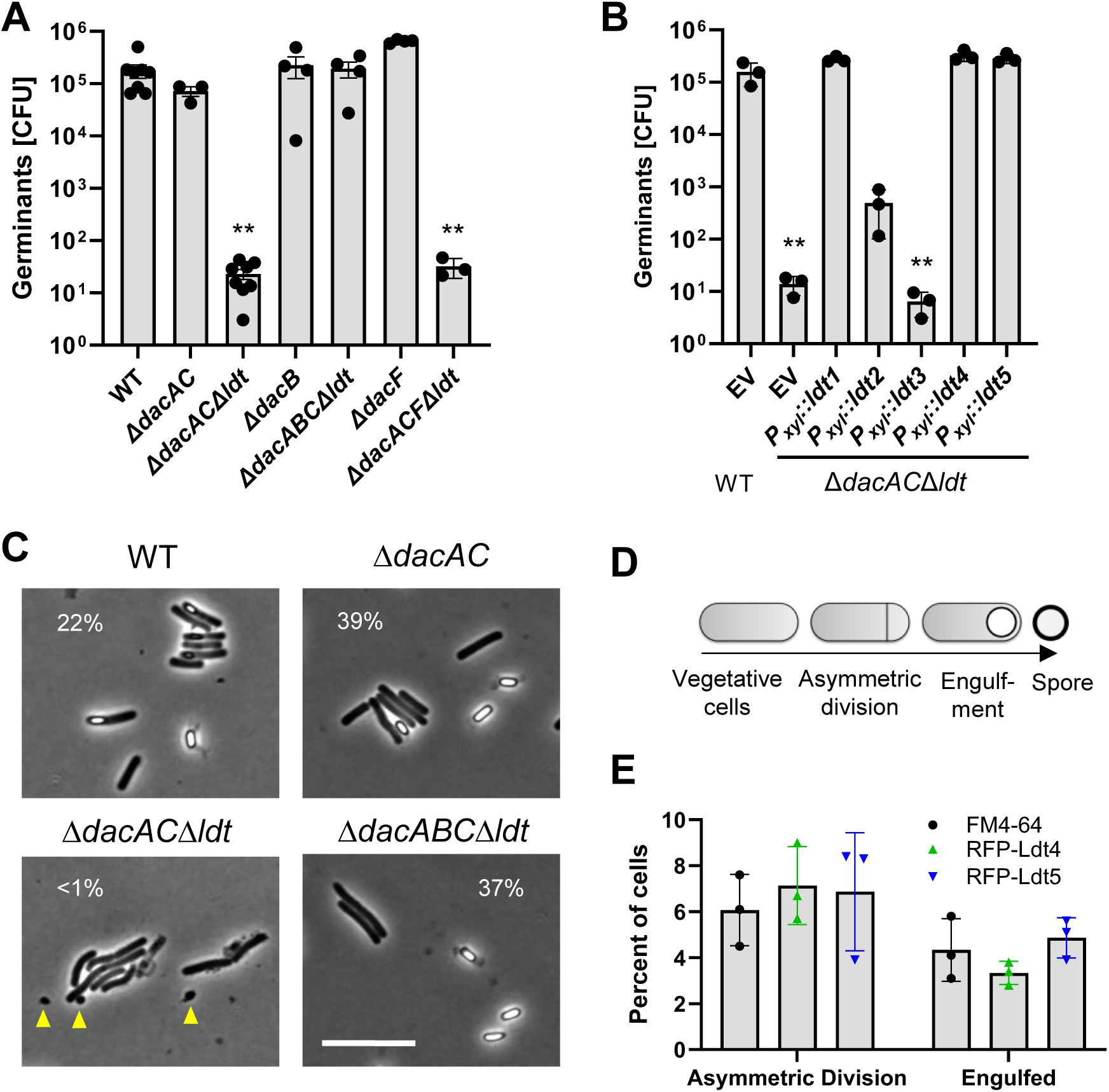
LDTs are required for sporulation. (A) The ability of WT and the indicated mutants to form spores was measured by determining the number of ethanol-resistant CFUs (i.e., spores) per million total CFUs (i.e., vegetative cells). Results are graphed as the mean and SD of 3-7 biological replicates performed on different days. To test for statistical significance, data were log-transformed and analyzed using one-way ANOVA followed by a Dunnett’s test comparing each strain to WT. **= p<0.0001 (B) Rescue of sporulation by complementation with the indicated LDTs produced from a P*_xyl_* expression vector. EV = empty vector. Statistical significance was calculated as in (A) comparing each strain to WT EV. **= p<0.0001 (C) Phase contrast micrographs of WT and mutants after 24 hours on sporulation plates. Percents indicate abundance of phase-bright spores to the total of vegetative cells and spores. Yellow carets indicate phase dark bodies that are probably immature spores. Size bar: 10 µm. (D) Visual representation of the stages of sporulation in *C. difficile*. (E) Quantification of localization of FM4-64 staining or RFP-LDT fusions to sites of asymmetric division or engulfment. Frequencies are low because most cells appeared to be vegetative. Data are graphed as the mean and SD from 3 biological replicates with a minimum of 102 cells each.

**FIG 7:**
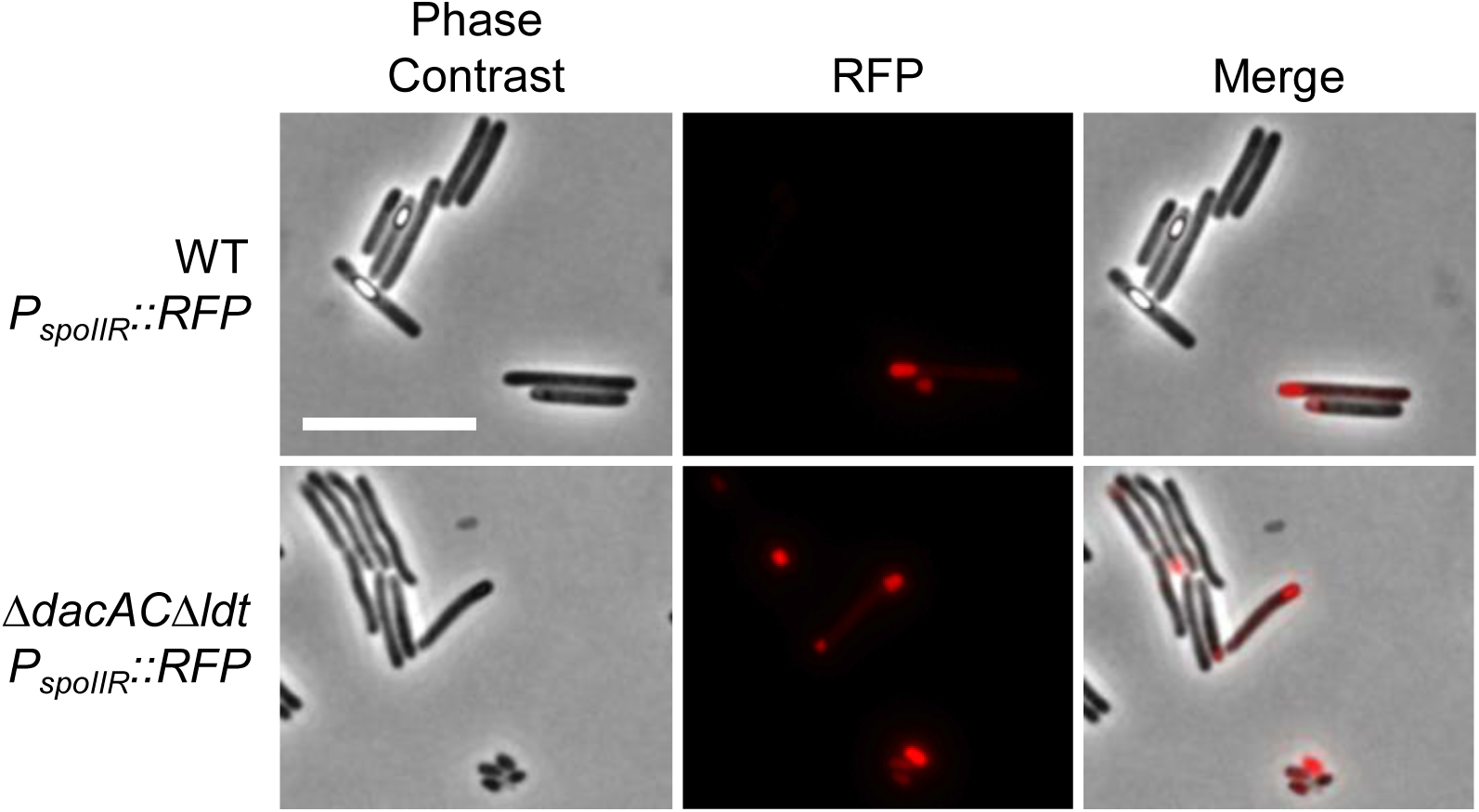
Sporulation is initiated in Δ*dacAC*Δ*ldt*. Microscopy of cells with a sporulation transcriptional reporter driving expression of RFP. Strains were grown on sporulation plates with Thi10 for 20 hours, scraped, resuspended, and fixed. Overnight incubation at 4°C allowed the chromophore to develop and cells were imaged by phase contrast and red fluorescence microscopy. *P_spoIIR_* is induced early during sporulation, so fluorescent cells have begun sporulating. Size bar: 10 µm.

### Ldt4 and Ldt5 localize to developing spores

For LDTs to crosslink spore PG, they must be present at the forespore during sporulation. To ascertain whether this is the case, WT cells with or without P*_xyl_* plasmids that express *rfp* fusions to *ldt4* and *ldt5* were plated on sporulation media containing a low level of xylose (0.02%) to produce the fusion proteins at native levels as determined by Western blotting (Fig. S3). Cells were recovered from plates after 24 hours, fixed, and visualized by fluorescence microscopy. Cells without an RFP fusion were stained with the DNA dye Hoechst 33342 and the membrane dye FM4-64 to determine the fraction of cells at various morphological stages in the sporulation process. Most cells were not sporulating, but approximately 6% were undergoing asymmetric division and 4% were undergoing engulfment (Fig. 6E, Fig. S4). In parallel assays, RFP-Ldt4 and RFP-Ldt5 were observed at asymmetric division sites in ∼6.5% of the cells and around the developing spore in ∼3.5% of the cells (Fig. 6E, Fig. S4). Thus, these LDTs are recruited to developing spores at frequencies consistent with a role in 3-3 crosslink formation.

### Loss of *dacB* restores sporulation to WT levels in Δ*dacAC*Δ*ldt1-5*

The above observations leave open the question of why deletion of *dacA* and *dacC* does not bypass the LDT requirement for sporulation, as it does for vegetative growth. We could envision several possible explanations. Rather than test them directly, we selected for suppressor mutations that restored sporulation to near WT levels in the Δ*dacAC*Δ*ldt* background. To this end, we enriched for mutants that can sporulate efficiently by subjecting the Δ*dacAC*Δ*ldt* strain to four rounds of sporulation followed by germination. We isolated two independent candidates that sporulated at a frequency about 1000-fold higher than the Δ*dacAC*Δ*ldt* parent strain but still about 10-fold lower than WT (Fig. S5). Whole genome sequencing revealed each suppressor strain had a mutation in *cdr20291_2048* (*dacB*), a DD-carboxypeptidase that is induced by SigF during sporulation [41]. One isolate had a frameshift at codon 11 and the other had a missense mutation that changes Asp209 to Tyr. Deletion of *dacB* in the Δ*dacAC*Δ*ldt* strain rescued sporulation to WT levels, confirming that loss of *dacB* suppresses the defect (Fig. 6A). In contrast, deletion of *dacB* in the WT background did not affect sporulation frequency (Fig. 6A). As expected, complementation of the Δ*dacABC*Δ*ldt* mutant with *dacB* reduced sporulation frequency to that of the Δ*dacAC*Δ*ldt* parent (Fig. S5).

In addition to *dacB*, there is a second sporulation-induced DD-CPase, *dacF*, which is expressed in the mother cell under the control of SigG [40]. To determine whether loss of *dacF* would rescue sporulation as observed for loss of *dacB*, we deleted *dacF* in WT and the Δ*dacAC*Δ*ldt* backgrounds. There was no rescue (Fig. 6A).

### Deletion of *dacB* increases crosslinking in spore PG

We hypothesized that the loss of DacB facilitated sporulation in the absence of LDTs by increasing 4-3 crosslinking mediated by PBPs. To test this hypothesis, we characterized the muropeptide profile of spores from WT, Δ*dacAC*, Δ*dacB*, and Δ*dacABC*Δ*ldt*. It was not possible to include Δ*dacAC*Δ*ldt* in this analysis as no spores could be purified. Representative elution profiles are shown in Fig. 8. Muropeptides were collected from Δ*dacB* samples to determine their structures by mass spectrometry. These identifications were extended to the other strains based on retention time in RP-HPLC.

**FIG 8:**
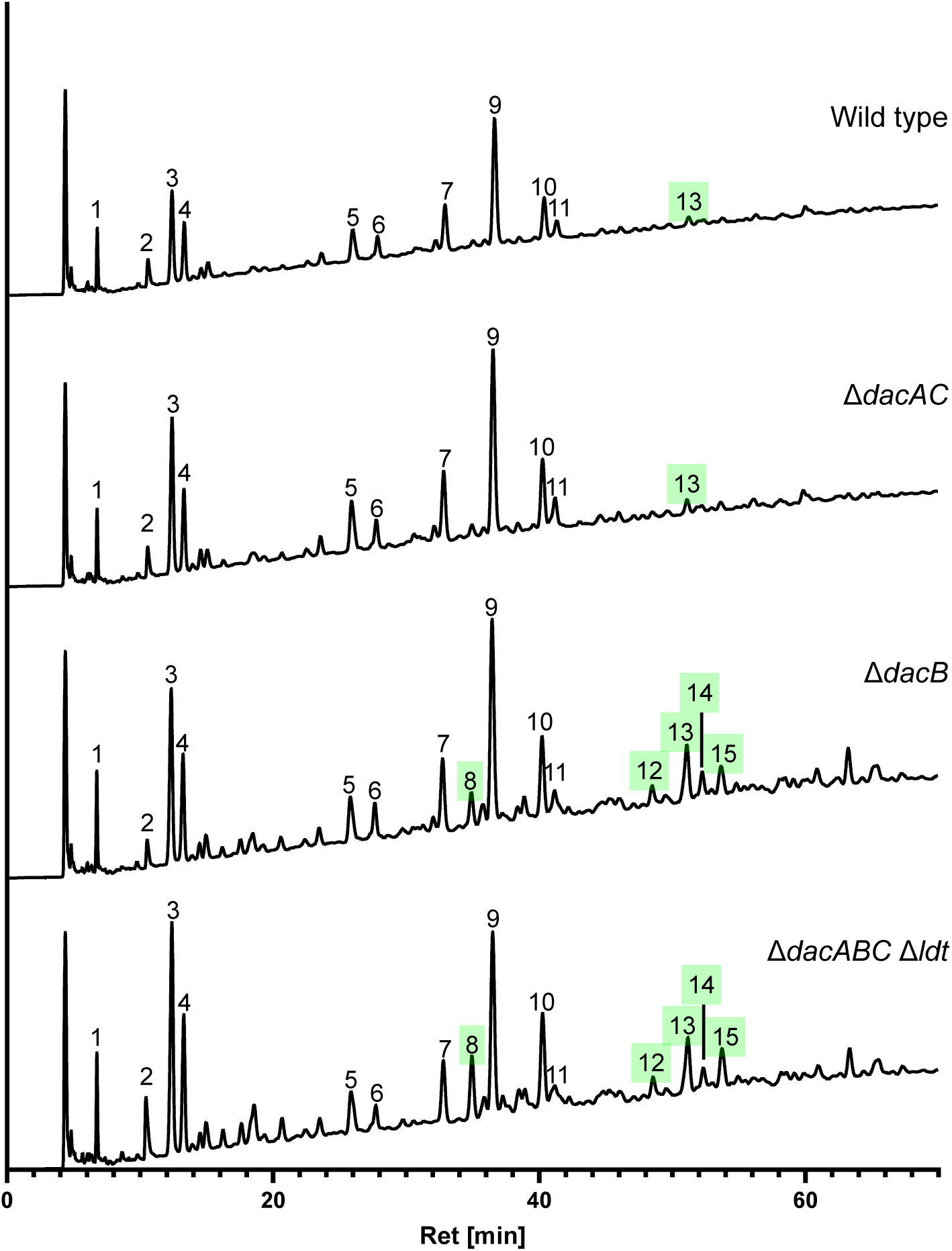
The absence of DacB increases 4-3 crosslinks in spore peptidoglycan. HPLC separation of muropeptides from the indicated strains. Peaks present at a relative abundance of ≥ 2% in the Δ*dacB* mutant are numbered. Peak identification and quantitation are summarized in Table 3. Peaks highlighted in green are 4-3 crosslinked dimers; 3-3 crosslinked dimers were not detected. Peaks 1 and 2 are not muropeptides. Chromatograms are representative of three biological replicates.

**Table 3.** Muropeptide quantitation of spore peptidoglycan.

| Peak<sup>1</sup> | Rt<sup>2</sup> [min] | m/z obs<br>[M+H]<sup>+</sup> | Muropeptide<sup>3</sup> | Cross-<br>link | Abundance [%]<sup>4</sup> |  |  |  |
| --- | --- | --- | --- | --- | --- | --- | --- | --- |
|  |  |  |  |  | WT | *ΔdacAC* | *ΔdacB* | *ΔdacABC*<br>*Δldt* |
| Monomers |  |  |  |  |  |  |  |  |
| 3 | 12.4 | 942.42 | DS-TP |  | 15.8 ± 0.9 | 16.3 ± 1.1 | 15.2 ± 0.3 | 19.2 ± 1.3 |
| 4 | 13.3 | 900.41 | DS-TP (deAc) |  | 8.9 ± 0.8 | 7.9 ± 0.7 | 8.2 ± 0.4 | 9.9 ± 0.8 |
| 5 | 25.9 | 1304.59 | Reduced TS-TP<br>(deAc)<sup>5</sup> |  | 7.5 ± 0.6 | 7.6 ± 0.5 | 5.9 ± 0.2 | 4.9 ± 0.5 |
| 6 | 27.7 | 1276.57 | TS-TP (2x deAc) |  | 4.1 ± 0.5 | 3.7 ± 0.3 | 3.5 ± 0.2 | 2.5 ± 0.5 |
| 7 | 32.8 | 1318.58 | TS-TP (2x deAc)l |  | 9.8 ± 0.7 | 10.1 ± 0.3 | 8.2 ± 0.4 | 6.5 ± 0.7 |
| 9 | 36.5 | 1318.58 | TS-TP (deAc) |  | 31.3 ± 2.3 | 30.7 ± 3.1 | 24.5 ± 1.3 | 20.4 ± 2.0 |
| 10 | 40.2 | 1360.58 | TS-TP |  | 10.2 ± 0.7 | 10.9 ± 0.5 | 9.8 ± 0.3 | 11.0 ± 0.7 |
| 11 | 41.2 | 1075.45 | TS-DiP (deAc) |  | 4.8 ± 0.5 | 5.0 ± 0.4 | 3.4 ± 0.2 | 3.4 ± 0.03 |
| Dimers |  |  |  |  |  |  |  |  |
| 8 | 34.9 | 1865.83 | DS-TP-TP-DS | 4-3 | 1.6 ± 0.3 | 1.8 ± 0.4 | 3.9 ± 0.3 | 6.3 ± 1.3 |
| 12 | 48.6 | 2241.98 | DS-TP-TP-TS<br>(deAc) | 4-3 | 1.2 ± 0.3 | 1.1 ± 0.2 | 2.5 ± 0.2 | 2.1 ± 0.2 |
| 13 | 51.2 | 2241.98 | DS-TP-TP-TS<br>(deAc) | 4-3 | 2.2 ± 0.2 | 2.4 ± 0.2 | 7.8 ± 0.7 | 6.8 ± 0.4 |
| 14 | 52.3 | 2199.97 | DS-TP-TP-TS<br>(2x deAc) | 4-3 | 1.1 ± 0.3 | 0.9 ± 0.2 | 3.1 ± 0.3 | 2.3 ± 0.2 |
| 15 | 53.7 | 2283.99 | DS-TP-TP-TS | 4-3 | 1.6 ± 0.3 | 1.6 ± 0.1 | 4.1 ± 0.3 | 4.5 ± 0.6 |

<sup>1</sup>Peaks 1 and 2 are not muropeptides
<sup>2</sup>Rt, retention time
<sup>3</sup>Abbreviations: DS: disaccharide (NAG-NAMol), TS: tetrasaccharide (NAG-MAL-NAG-NAMol), DiP: dipeptide (L-Ala-D-Glu), TP: tetrapeptide (L-Ala-D-Glu-mDAP-D-Ala), deAc: deacetylation of NAG.
<sup>4</sup>Abundance is reported as the mean ± SD from three biological replicates and was calculated as the ratio of the peak area to the sum of the areas of all the muropeptide peaks identified in the table.
<sup>5</sup>Here "reduced" refers to reduction of the MAL lactam by borohydride (NAG-MAL<sup>red</sup>-NAG-NAMol).

The deduced structures of 13 muropeptides are presented in Table 3, along with their abundance as a percentage of total analyzed muropeptides. Of note, and in contrast to vegetative PG, eight of the 13 muropeptides contain a tetrasaccharide moiety. This is consistent with previous reports [17, 35, 36] and is explained by conversion of ∼25% of the NAM residues to muramic-δ-lactam, a hallmark of spore cortex PG [33]. Another hallmark of spore cortex PG apparent in our samples is low levels of crosslinking [33]. In accordance with earlier studies, ∼92% of total muropeptides in WT samples were monomers, ∼8% were dimers, and trimers were not detected [17, 35, 36]. Surprisingly, however, all of the dimers were 4-3 crosslinked. We had anticipated predominantly 3-3 crosslinking based on a recent report [17]. The absence of 3-3 crosslinked dimers in our analyses likely reflects their low abundance as even the most prominent 3-3 crosslinked dimer in the previous study constituted only ∼1.7% of total muropeptides. In contrast, the smallest peak that we collected and characterized was peak 12 and amounted to 2.5% of the total muropeptide in Δ*dacB* samples (Fig. 8, Table 3). Regarding the 4-3 crosslinked dimers, it can be excluded that they are due to contamination with vegetative PG because four of them contain a tetrasaccharide (peaks 12-15) diagnostic of cortex PG. Moreover, two previous analyses of *C. difficile* spore PG reported dimeric muropeptides with two tetrapeptides, which can form a 4-3 crosslink but not a 3-3 crosslink [35, 36].

Turning now to the various mutants, there were no significant differences between the muropeptide composition of WT and Δ*dacAC* spores. But when *dacB* was deleted, 4-3 crosslinked dimers more than doubled to 22% of the total (peaks 8, 12-15). This is consistent with our hypothesis and implies that the loss of DacB increases the pool of PG pentapeptides, which in turn promotes 4-3 crosslinking by PBPs required for sporulation [42]. Combining Δ*dacB* with deletions of *dacA*, *dacC* and the five *ldt* genes did not result in any noteworthy additional changes to the muropeptide profile.

## Discussion

The high fraction of 3-3 crosslinks in *C. difficile* PG could be explained if most of the PG precursor is converted from pentapeptide to tetrapeptide by DD-CPases, as recently suggested by Oliveira Pavia et al. based on their observation that growth in the presence of a sublethal concentration of β-lactams increases the abundance of 4-3 crosslinks at the expense of 3-3 crosslinks [17]. Here we confirmed and extended that finding by identifying DacA and DacC as the DD-CPases responsible for supplying LDTs with tetrapeptides during vegetative growth. A *dacAC* double deletion decreased 3-3 crosslinks, increased 4-3 crosslinks and bypassed the normal requirement for LDTs for viability. Thus, we were able to delete all five *ldt* genes in a Δ*dacAC* background to produce a derivative of *C. difficile* that grows with an exclusively 4-3 crosslinked PG sacculus. This derivative was not just viable, it was healthy. It grew at the same rate as WT, with only subtle morphological defects, no increased sensitivity to β-lactams, and only modestly increased sensitivity to glycopeptide antibiotics. The most striking phenotypic defect was a 3- to 4-log decrease in viable spores, which, however, was overcome by deletion of a sporulation-induced DD-CPase, *dacB*.

Besides identifying the source of tetrapeptides for 3-3 crosslinking, our findings reveal that LDTs are required for viability because *C. difficile* has intrinsically high DD-CPase activity rather than because of anything special about 3-3 as opposed to 4-3 crosslinks. LDTs are not essential components of the divisome or the elongasome, although they may be nonessential components of these complexes. Moreover, conversion of pentapeptide precursors to tetrapeptides limits PBP activity and explains our previous observation that *C. difficile* does not increase 4-3 crosslinks to compensate for loss of 3-3 crosslinks when cells are depleted of LDTs [12]. In other words, LDTs and PBPs are not in competition, and conversion of pentapeptides to tetrapeptides by DD-CPases effectively sidelines the PBPs even when LDTs are absent.

Based on these findings, we propose a model for how PBPs and LDTs work together in *C. difficile* (Fig. 9). In this model, *C. difficile* synthesizes lipid II with a pentapeptide sidechain as is the case in other bacteria. The precursor is transported across the membrane to the cell’s exterior, where DD-CPases convert much (but not all) of it to tetrapeptide. Bifunctional PBPs and SEDS-PBP complexes utilize the mixed pool of precursors to polymerize glycan strands that are sparsely crosslinked due to the inability of PBPs to use tetrapeptides as acyl donors. LDTs provide additional crosslinks to stabilize the sacculus against osmotic stress. The LDTs might function autonomously or in complexes with the PBPs. However, even if the latter is the case, the LDTs do not have any essential roles beyond crosslinking since PBPs are fully capable of building the sacculus (and spore cortex) in the complete absence of LDTs, provided sufficient pentapeptides precursors are available.

**FIG 9:**
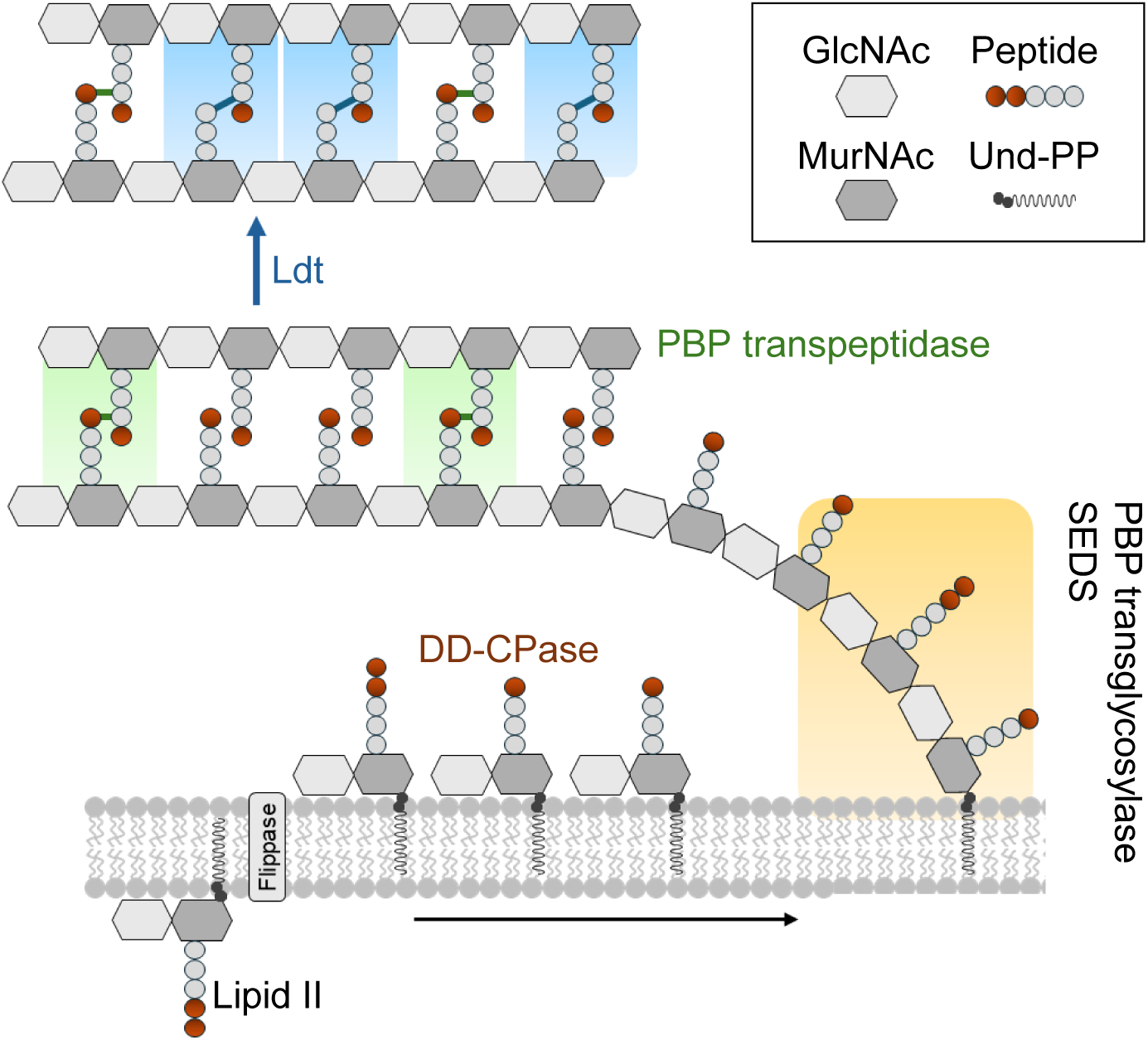
Model for how PBPs and LDTs work together in *C. difficile.* Lipid II is assembled on the cytoplasmic face of the cell membrane. An unidentified flippase transports lipid II outside the cell, where DD-CPases convert many of the pentapeptide sidechains to tetrapeptides. The mixed pool of precursors is polymerized and crosslinked by SEDS proteins and PBPs, but crosslinks are sparse owing to the low abundance of pentapeptides. LDTs subsequently introduce 3-3 crosslinks needed for the sacculus to withstand turgor pressure.

The dependency of LDTs on multiple DD-CPases has implications for the potential to exploit LDTs as targets for antibiotics that kill *C. difficile* selectively. Such drugs will be of little use if it is easy for *C. difficile* to become resistant. The fact that three DD-CPase genes had to be deleted to bypass the requirement for LDTs for growth and sporulation suggests it would be difficult to evolve resistance by this pathway. Other pathways no doubt exist, such as remodeling LDT active sites to exclude the antibiotic, but evaluating the frequency and efficacy of those mechanisms will not be possible until suitable drugs are in hand.

The dependency of LDTs on DD-CPases also has implications for β-lactam resistance. *C. difficile* is remarkably resistant to one class of β-lactams, the cephalosporins. These drugs inactivate PBPs but not LDTs. Cephalosporin resistance might therefore be explained by the ability of LDTs to sustain PG crosslinking when PBPs are no longer active. However, that scenario predicts that abrogating the LD-transpeptidation pathway should sensitize *C. difficile* to cephalosporins, which is not what we observed—the MIC for cefoxitin (a cephalosporin) was the same in WT and the Δ*dacAC* Δ*ldt* mutant. Moreover, the MICs for ampicillin (a penicillin) and meropenem (a penem) were also unchanged.

These counter-intuitive observations can be explained in at least two ways, which are not mutually exclusive. First, PBP1 and PBP2 are essential in *C. difficile*, so even small reductions in their activity might not be tolerated, especially considering that only ∼30% of the crosslinks are 4-3 crosslinks to begin with [9, 11, 13, 43]. In support of this hypothesis, the MICs for multiple β-lactams correlate with their affinities for PBP1 and PBP2 [44]. Alternatively, the notion that LDTs can sustain crosslinking in the face of β-lactam stress might not be true in *C. difficile*. LDTs require tetrapeptide acyl donors generated by β-lactam-sensitive DD-CPases, so inactivation of these would inhibit the LD-transpeptidation pathway even though the LDTs are not acylated. Consistent with this hypothesis, sublethal doses of β-lactams skew crosslinking from 3-3 to 4-3 [17]. If the primary targets of these antibiotics were PBP1 and PBP2, 4-3 crosslinking should have gone down before 3-3 crosslinking. In summary, it appears that β-lactams inhibit 4-3 crosslinking directly by acylation of PBP1 and PBP2, and 3-3 crosslinking indirectly by depriving the LDTs of tetrapeptide acyl donors. The weaker link may be the LD-transpeptidation pathway [17].

The inability of LDTs to confer resistance to β-lactams in *C. difficile* contrasts with lab-derived mutants of *Enterococcus faecium* and *E. coli* that are highly β-lactam resistant because they rely exclusively on LDTs to crosslink PG [15, 45]. The latter organisms use β-lactam insensitive DD-CPases to generate tetrapeptides, either a cytoplasmic metallo-CPase in the case of *E. faecium* or a periplasmic serine-type CPase with low affinity for β-lactams in the case of *E. coli*.

Vancomycin is the frontline treatment for *C. difficile* infections, but resistance is on the rise [46, 47]. Vancomycin binds to the D-Ala^4^-D-Ala^5^ termini of PG precursors, so it is somewhat surprising that *C. difficile* is vancomycin sensitive despite relying primarily on LDTs for crosslinking. There are at least two potential explanations. One is that vancomycin inhibits not only transpeptidation but also glycan strand polymerization. The latter activity cannot be substituted by LDTs. The other is that vancomycin prevents DD-CPases from converting pentapeptides to tetrapeptides and thus inhibits 3-3 crosslink formation indirectly. We observed that LDTs make modest contributions to *C. difficile’s* intrinsic resistance to vancomycin and related glycopeptide antibiotics.

One factor that makes *C. difficile* infections difficult to eradicate is that the organism forms metabolically dormant spores. Spores are surrounded by a complex, multilayered envelope, including two layers of PG. These are the germ cell wall and the cortex. The germ cell wall is structurally similar to vegetative PG, while the cortex is thick, highly modified, and sparsely crosslinked. A specialized PG synthase, consisting of the SEDS family glycosyltransferase SpoVE and its cognate transpeptidase SpoVD (a PBP), is induced during sporulation and has been shown by several groups to be essential for cortex synthesis [9, 40, 48]. Whether LDTs are also essential for sporulation was not yet known but seemed likely given that spore PG contains both 3-3 and 4-3 crosslinks [17, 35, 36]. The fact that LDTs are essential for vegetative growth has until now prevented testing their importance during sporulation.

We observed an approximately 6000-fold reduction in ethanol-resistant spores in the Δ*dacAC*Δ*ldt* mutant compared to WT. Follow-up experiments revealed that development goes awry after asymmetric division and results in the production of immature, phase-dark spores. Selecting for suppressor mutations that enabled the Δ*dacAC*Δ*ldt* mutant to sporulate revealed the LDT requirement can be bypassed by loss of *dacB*, which encodes a DD-CPase induced in the forespore early during its development. Analysis of spore PG from Δ*dacB* showed an increase in 4-3 crosslinks compared to WT, confirming that loss of this DD-CPase provided PBPs with more substrate. Interestingly, deletion of another sporulation-induced DD-CPase, *dacF*, had no effect on LDT essentiality for sporulation. We hypothesize that the timing of when the DD-CPases are expressed determines their involvement in LDT importance during sporulation, as *dacB* is expressed early under control of σ^F^, whereas *dacF* is expressed later under control of σ^G^. Further studies will be needed to determine whether LDTs are required for synthesis of the germ wall, the cortex, or both. However, two observations implicate LDTs in crosslinking the germ cell wall during engulfment. First, loss of σ^F^-controlled DacB but not σ^G^-controlled DacF eliminates the need for LDTs during sporulation. Second, microscopy suggests the forespore and mother cells lose integrity when DacB is active in the absence of LDTs.

## Materials and Methods

### Strains, media, and growth conditions

Bacterial strains are listed in Table 4. *C. difficile* strains used in this study were all derived from R20291 [49]. *C. difficile* was grown in tryptone-yeast (TY) media, supplemented as needed with thiamphenicol at 10 μg/mL (Thi_10_), kanamycin at 50 μg/mL, or cefoxitin at 8 μg/mL. Anhydrous tetracycline (aTet) was used to induce genes under P*_tet_* control (cat. number 37919, Sigma Aldrich, Burlington, MA). TY media consisted of 3% tryptone, 2% yeast extract, and 2% agar (for solid media). For conjugation plates brain heart infusion (BHI, Bacto) solid media was used. BHI media consisted of 3.7% BHI and 2% agar. BHIS media was BHI supplemented with 5% yeast extract. *C. difficile* strains were grown at 37°C in an anaerobic chamber (Coy Laboratory Products, Grass Lake, MI) in an atmosphere of about 2% H_2_, 5% CO_2_, and 93% N_2_. Growth in broth in glass tubes was monitored at OD_600_ with a WPA Biowave CO8000 Cell Density Meter. Continuous growth curves were monitored in a Tecan Sunrise plate reader in flat-bottom 96-well plates with 200 µL of culture.

**Table 4:** Strains used in this study.

| Strain Number | Genotype | Plasmid Features | Reference or Source |
| --- | --- | --- | --- |
| R20291 | Wild type strain from UK outbreak (ribotype 027) |  | [49] |
| KB464 | R20291/pBZ101 | *P<sub>xyl</sub>* EV | [12] |
| KB449 | R20291/pHL14 | *P<sub>xyl</sub>::mCherry<sup>Opt</sup>-ldt4* | [13] |
| KB455 | R20291/pHL15 | *P<sub>xyl</sub>::mCherry<sup>Opt</sup>-ldt5* | [13] |
| KB547 | R20291 Δldt1-3Δldt4 *P<sub>tet</sub>::ldt5* | | [12] |
| KB650 | R20291 Δldt1-4ΔdacA *P<sub>tet</sub>::ldt5* | | This study |
| KB661 | R20291 Δldt1-4ΔdacC *P<sub>tet</sub>::ldt5* | | This study |
| KB803 | R20291 ΔdacAΔdacC | | This study |
| KB834 | R20291 Δldt1-4ΔdacD *P<sub>tet</sub>::ldt5* | | This study |
| KB856 | R20291 Δldt1-4ΔdacCΔdacD *P<sub>tet</sub>::ldt5* | | This study |
| KB863 | R20291 ΔdacAΔdacCΔdacD | | This study |
| KB874 | R20291 Δldt1-4ΔdacAΔdacD *P<sub>tet</sub>::ldt5* | | This study |
| KB880 | R20291/pRAN739 | *P<sub>spolIR</sub>::mCherry<sup>opt</sup>* | This study |
| KB929 | R20291 ΔdacF | | This study |
| KB955 | R20291 ΔdacB | | This study |
| K1069 | R20291 Δldt1-4ΔdacAΔdacCΔdacD *P<sub>tet</sub>::ldt5* | | This study |
| KB1001 | R20291 Δldt1-4ΔdacAΔdacC *P<sub>tet</sub>::ldt5* | | This study |
| KB1008 | R20291 Δldt1-5ΔdacAΔdacC | | This study |
| KB1018 | R20291 Δldt1-5ΔdacAΔdacC/pBZ101 | *P<sub>xyl</sub>* EV | This study |
| KB1019 | R20291 Δldt1-5ΔdacAΔdacC/pKB25 | *P<sub>xyl</sub>::ldt1* | This study |
| KB1020 | R20291 Δldt1-5ΔdacAΔdacC/pCE938 | *P<sub>xyl</sub>::ldt2* | This study |
| KB1021 | R20291 Δldt1-5ΔdacAΔdacC/pCE983 | *P<sub>xyl</sub>::ldt3* | This study |
| KB1022 | R20291 Δldt1-5ΔdacAΔdacC/pCE1175 | *P<sub>xyl</sub>::ldt4* | This study |
| KB1023 | R20291 Δldt1-5ΔdacAΔdacC/pCE1176 | *P<sub>xyl</sub>::ldt5* | This study |
| KB1024 | R20291 Δldt1-5ΔdacAΔdacBΔdacC | | This study |
| KB1052 | R20291 Δldt1-5ΔdacAΔdacCΔdacF | | This study |
| KB1054 | R20291/pKB157 | *P<sub>dacB</sub>::dacB* | This study |
| KB1056 | R20291 Δldt1-5ΔdacAΔdacBΔdacC/pBZ101 | *P<sub>xyl</sub>* EV | This study |
| KB1057 | R20291 Δldt1-5ΔdacAΔdacBΔdacC/pKB157 | *P<sub>dacB</sub>::dacB* | This study |
| KB1058 | R20291 Δldt1-4ΔdacAΔdacC *P<sub>tet</sub>::ldt5*/pBZ101 | *P<sub>xyl</sub>* EV | This study |
| KB1059 | R20291 Δldt1-4ΔdacAΔdacC *P<sub>tet</sub>::ldt5*/pKB92 | *P<sub>xyl</sub>::dacA* | This study |
| KB1060 | R20291 Δldt1-4ΔdacAΔdacC *P<sub>tet</sub>::ldt5*/pKB158 | *P<sub>dacC</sub>::dacC* | This study |
| KB1069 | R20291 Δldt1-4ΔdacAΔdacCΔdacD *P<sub>tet</sub>::ldt5*/pKB158 | *P<sub>dacC</sub>::dacC* | This study |
| KB1103 | R20291 Δldt1-5ΔdacAΔdacCΔdacD | | This study |
| KB1104 | R20291 Δldt1-5ΔdacAΔdacC/pRAN739 | *P<sub>spolIR</sub>::mCherry<sup>opt</sup>* | This study |
| PB102 | R20291 Δldt1-5ΔdacAΔdacCΔdacD *dacB<sup>D209T</sup>* | | This study |
| PB105 | R20291 Δldt1-5ΔdacAΔdacCΔdacD *dacB<sup>I11fs</sup>* | | This study |

*Escherichia coli* strains were grown in LB media at 37°C with chloramphenicol at 10 μg/mL or ampicillin at 100 μg/mL as necessary. LB media contained 1% tryptone, 0.5% yeast extract, 0.5% NaCl, and 1.5% agar (for solid media).

### Plasmid and bacterial strain construction

Plasmids are listed in Supplemental Table S1 and were constructed by isothermal assembly with reagents from New England Biolabs (NEB, Ipswich, MA). Regions that were constructed by PCR were verified by DNA sequencing. The oligonucleotide primers used in this study were synthesized by Integrated DNA Technologies (Coralville, IA) and are listed in Table S2. All plasmids were propagated using OmniMax 2-T1R as the cloning host. Plasmids were moved into *C. difficile* by first transforming into HB101/pRK24 [50, 51] then conjugating into *C. difficile* [52]. CRISPR editing plasmids were designed as previously described [53] with a single guide RNA against the target gene and homology regions to repair the double-stranded break caused by the Cas9 nuclease. Mutagenesis was induced by plating R20291 harboring an editing plasmid onto TY Thi_10_ with 1% xylose to induce expression of *cas9*. Typical plating efficiency was around 10^-4^ but varied by construct. Survivor colonies were restruck once on TY with 1% xylose and subsequently maintained by serial passage on TY until the plasmid was lost as evidenced by sensitivity to thiamphenicol. Successful mutagenesis was confirmed by PCR with Q5 or Taq DNA polymerase (NEB).

### Viability plating

Viability was tested by spot titer assays. For this, a 10-fold dilution series was prepared from overnight cultures, and 5 μL of each dilution were spotted onto the appropriate plates, which were incubated at 37°C overnight before imaging.

### Microscopy

Cells were immobilized using thin agarose pads (1%). Phase-contrast and fluorescence micrographs were recorded on an Olympus BX60 microscope equipped with a 100× UPlanXApo objective (numerical aperture, 1.45). Micrographs were captured with a Hamamatsu Orca Flash 4.0 V2+ complementary metal oxide semiconductor (CMOS) camera. Excitation light was generated with an X-Cite XYLIS LED light source. Membranes were stained with the lipophilic dye FM4-64 (Life Technologies, Carlsbad, CA) at 1 μg/mL. Cells were imaged immediately without washing. Red fluorescence was detected with the Chroma filter set 49008 ET (ex 538-582 nm excitation filter, em 590-667 nm emission). DNA was stained with 10 μg/mL Hoechst 33342 (Invitrogen, Waltham, MA). Cells were incubated at room temperature for 15 min and imaged without washing. Blue fluorescence was detected with the Chroma filter Set 49000 ET (ex 326-375 nm, em 436-485 nm). MicrobeJ was used to measure cell length, width, and sinuosity [54].

### Time-lapse microscopy

Growing cells were imaged anaerobically as described [24] with the following modifications. The day before the experiment, a 125 µL Gene Frame (Thermo Scientific, Waltham, MA) attached to a glass microscope slide, 2% molten agarose (RPI Laboratory Products, Swedesboro, MA), and 2x TY were brought into the anaerobic chamber. The agarose was maintained at 68°C to keep it molten. The next day, the 2x TY was warmed to 68°C, then mixed 1:1 with the 2% agarose. An excess of this mixture (500uL) was added into the Gene Frame well and covered with a second microscope slide. After letting the agarose solidify for 5 minutes, the upper microscope slide was carefully slid off the agarose pad. Bacterial samples (5 µl) were spotted onto the agarose, allowed to dry (∼5 min), and the slide was sealed with a 24 x 50 mm glass coverslip. Slides were mounted on a heated (37°C) microscope stage and imaged every 5 minutes using an inverted Nikon Ti2 Eclipse microscope as described [55]. Movies and montages were generated using Nikon NIS-Elements AR software.

### Sporulation

Sporulation frequencies were determined using an ethanol resistance assay to quantify the ratio of ethanol-resistant spores to total viable cells as previously described [56]. Briefly, cells were incubated on 70/30 sporulation plates for 24hrs. Cells and spores were scraped from the plates into approximately 5 mL of BHIS to an OD_600_ of 1. To determine total viable cells, dilutions were plated on BHIS agar. To determine the spore count, an aliquot was treated with ethanol (50%) for 15 minutes to eliminate vegetative cells, then serially diluted and plated on BHIS agar with 0.1% taurocholate. The plates were incubated anaerobically for 24 hours, and CFUs were counted to calculate sporulation frequency as a percentage of total viable cells.

The selection for mutants in the Δ*dacAC*Δ*ldt* background used repeated sporulation-germination cycles. An overnight culture of Δ*dacAC*Δ*ldt* grown in BHIS with 0.2% D-fructose and 0.1% taurocholate was subcultured 1:100 into the same media. When it grew to an OD_600_ of 0.5, 250 µl were spread on a 70/30 sporulation plate. After 24 h, spores were isolated and vegetative cells killed with ethanol (50%), then plated on BHIS agar with 0.1% taurocholate. The resulting colonies were pooled and grown overnight in liquid BHIS with fructose and taurocholate. The next day, the culture was diluted 1:100 in the same media, grown to OD_600_ of 0.5 and plated again on 70/30 sporulation media. This process was repeated four times until sporulation efficiency was within 10-fold of WT, at which point a single colony was isolated, and DNA was extracted for whole genome sequencing as described below.

To visualize localization of RFP-LDT fusions during sporulation, R20291 harboring P*_xyl_*::*RFP-ldt* fusion plasmids was inoculated onto sporulation plates with Thi_10_ and 0.02% xylose. After 24 h cells and spores were removed by repeated scraping with an inoculating loop, suspended in approximately 5 mL BHIS to an OD_600_ of 1. Then 100 µL sample was fixed as described [12]. Fixed cells were maintained at 4°C overnight in aerobic conditions for the chromophore to develop. After adding Hoechst 33342 dye to stain DNA, cells were imaged as described above. To determine the frequency of cells in different morphological stages of sporulation, R20291 was recovered from sporulation plates and stained with FM4-64 and Hoechst 33342.

### Whole genome sequencing

DNA was extracted from 5 mL overnight cultures as follows. Cells were harvested by repeated centrifugation in a 1.5 mL microfuge tube. The cell pellet was resuspended in 300 µL Thermopol buffer (NEB) and 4 units of Proteinase K (NEB), then incubated at 37°C until the suspension cleared, at least 1 hour. Lysed cells were added to 500 µL of TRIS buffered phenol/choroform/isoamyl alcohol 25:24:1 (BP17521, Fisher Scientific, Pittsburgh, PA) and mixed by vortexing until emulsified. Samples were centrifuged at 16,000 rcf for 5 minutes and the aqueous layer was carefully pipetted into a clean microfuge tube. To precipitate the DNA, 40 µL of 3M sodium acetate and 1 mL of ice cold 95% ethanol were added to the extracted DNA. DNA was pelleted by centrifugation at 16,000 rcf for 1 min. The pellet was washed with 1mL of 70% ethanol and air dried for at least 15 minutes. DNA was resuspended in 100 µL of water. DNA sequencing was performed by SeqCenter (Pittsburgh, PA) using 150-bp paired-end reads on Illumina NextSeq 2000. SNP analysis was performed at the BV-BRC (formerly PATRIC) website [57, 58]. The sequences were aligned to the R20291 genome (reference genome 645463.57) using BWA-mem [59] and SNPs identified with FreeBayes [60].

### Peptidoglycan purification from vegetative cells

Peptidoglycan (PG) from vegetative cells was purified as previously published [12, 61]. Briefly, cell pellets from a 100 mL culture grown in TY to OD_600_ ∼ 0.3 were boiled in SDS, then stripped of unwanted biological material through treatment with DNAse, RNAse, trypsin, hydrofluoric acid, and alkaline phosphatase. PG was thoroughly washed with water and stored at −20°C.

### Peptidoglycan purification from spores

PG purification from spores was adapted from published procedures [36, 62]. *C. difficile* strains were grown in TY to an OD_600_ of about 0.5 and 250 µL were spread on 70:30 sporulation plates [56]. Plates were incubated at 37°C for 5 days. Both cells and spores were scraped from the plates with an inoculating loop and suspended in 5 mL of cold water. Suspensions were maintained at 4°C overnight. Suspensions were carefully layered on top of 5 mL 50% HistoDenz (cat. number D2158-100G, Sigma Aldrich) solution (w/v) in a 15 mL plastic conical tube and centrifuged for 10 min at 7,000 rcf in a fixed-angle rotor to pellet the spores but not cell debris. Supernatants were carefully aspirated, and the spores washed with 1 mL water three times. The spore pellets were then resuspended in 500 µL of water and dilutions were made to measure the OD_600_. For each sample, a volume sufficient to contain ∼30 OD_600_ units was pelleted in a microfuge tube. Pellets were taken up in 1 mL 50 mM Tris-HCL, pH 7.5, 1% SDS, 50 mM DTT and boiled for 20 min, after which spore PG was recovered by centrifugation in a microfuge at 16000 rcf for one minute. Pellets were washed repeatedly by centrifugation and resuspension in water until free of SDS as determined using methylene blue [63]. The resulting pellets were resuspended in 1 mL 20 mM Tris-HCL, pH 8, 10 mM CaCl_2_, 0.1 mg/mL trypsin (LS003740, Worthington Biochemical, Lakewood, NJ) and incubated at 37°C for 20 hours. Trypsin was inactivated by adjusting the sample to 1% SDS and heating to 100°C for 15 min. PG was thoroughly washed with water to remove SDS and stored at −20°C.

### Muropeptide analysis

PG purified from vegetative cells or spores was digested with mutanolysin (M9901-10KU, Sigma Aldrich), then reduced with NaBH_4_, and resulting muropeptides separated by HPLC as described [12]. RP-HPLC with a NaPO_4_ buffer-methanol gradient yielded the elution profiles shown in Figs. 4 and 8. For mass spectrometry analysis, select peaks were collected and purified by RP-HPLC with a volatile buffer containing 0.1% trifluoroacetic acid and a gradient to 30% acetonitrile. Peaks were collected, lyophilized and LC-MS analyses were performed as described [12, 64].

### Western Blot

For the TcdA and TcdB Western blots, strains were grown in TY for 24 hours and the entire culture volume (10 mL) was centrifuged at 7,000 rcf for 10 minutes to remove the cells. The supernatant was collected, concentrated to a volume of approximately 250 µL using 100K centrifugal filters (Amicon), and mixed 1:1 with 2X Laemmli buffer. After heating at 95 °C for 10-15 min, 20 μL samples were electrophoresed on 10% sodium dodecyl sulfate-polyacrylamide gels (TGX gel, BioRad, Hercules, CA). Proteins were transferred to nitrocellulose and blots were developed using standard procedures. Primary antibodies against TcdA and TcdB were ACDTA and ACDTB, respectively (Fisher Scientific). These antibodies were used at a dilution of 1:5,000 and incubated with blots for 12 hours. Secondary antibody (IRDye 800 CW donkey anti-chicken, Li-COR, Lincoln, NE) was used at 1:10,000 and incubated for 1 hour. Blots were visualized with an Azure Biosystems Sapphire Biomolecular Imager.

Rabbit polyclonal antiserum against the soluble extracellular domains of Ldt4 and Ldt5 was raised by ProSci (San Diego, CA). The protein antigens were purified as described [12]. Primary antisera were diluted 1:10,000 and incubated with blots for 12 hours. Secondary antibody (IRDye 680LT goat anti-rabbit antibody, LI-COR, Lincoln, NE) was used at 1:10,000 for 1 hr. To verify antibody specificity, WT and Δ*ldt4* or Δ*ldt5* strains were grown in 3 mL TY to OD_600_ ∼ 0.85. Cells were harvested by centrifugation, resuspended in 300 μL 1X Laemmli buffer, and sonicated (Branson Sonifier 450, microtip, output 3, two cycles of 15 pulses) to generate whole cell lysates. Subsequent processing was as described above. Comparison of the abundance of RFP-Ldt4 and RFP-Ldt5 to the respective native proteins in sporulating cells used aliquots of the same samples that were harvested from sporulation plates for microscopy described above but without fixation. Specifically, 3 mL samples at an OD_600_ = 1.0 were centrifuged. The supernatants were discarded and the pellets containing cells and spores were taken up in 300 µL 1X Laemmli buffer. Subsequent steps were as described for TcdA and TcdB Westerns except for the use of different antibodies.

### Minimum inhibitory concentration assays

Susceptibility to antibiotics was determined in biological duplicate on two separate days as described [65].

## Data, Material, and Software Availability

The whole-genome sequencing data of Δ*dacAC*, Δ*ldt1-5*Δ*dacA*Δ*dacC*, Δ*ldt1-5*Δ*dacA*Δ*dacC*Δ*dacD dacB^D209T^*, and Δ*ldt1-5*Δ*dacA*Δ*dacC*Δ*dacD dacB^I11fs^* mutants were deposited under the BioProject ID PRJNA1482568.

## Acknowledgements

This work was supported by the National Institutes of Health grant R01 A1104821 (D.S.W. and C.D.E.) from the National Institute of Allergy and Infectious Diseases. Funding support to D.L.P. was through the National Institute of General Medical Sciences (GM138630). The LCMS work was funded by GlycoMIP, a National Science Foundation Materials Innovation Platform funded through Cooperative Agreement DMR-1933525. We thank Richard F. Helm for LC-MS analyses and members of the Ellermeier and Weiss laboratories for helpful discussions.

